# Pharmacologic rescue of circadian β-cell failure through P2Y1 purinergic receptor identified by small-molecule screen

**DOI:** 10.1101/2021.11.05.467499

**Authors:** Biliana Marcheva, Benjamin J. Weidemann, Akihiko Taguchi, Mark Perelis, Kathryn Moynihan Ramsey, Marsha V. Newman, Yumiko Kobayashi, Chiaki Omura, Jocelyn E. Manning Fox, Haopeng Lin, Patrick E. MacDonald, Joseph Bass

## Abstract

The mammalian circadian clock drives daily oscillations in physiology and behavior through an autoregulatory transcription feedback loop present in central and peripheral cells. Ablation of the core clock within the endocrine pancreas of adult animals impairs the transcription and splicing of genes involved in hormone exocytosis and causes hypoinsulinemic diabetes. However, identification of druggable proteins and pathways to ameliorate the burden of circadian metabolic disease remains a challenge. Here, we generated β cells expressing a nano-luciferase reporter within the proinsulin polypeptide to screen 2,640 pharmacologically-active compounds and identify insulinotropic molecules that bypass the secretory defect in clock mutant β cells. We validated lead compounds in primary mouse islets and identified known modulators of ligand-gated ion channels and G-protein coupled receptors, including the antihelmintic ivermectin. Single-cell electrophysiology in circadian mutant mouse and human cadaveric islets validated ivermectin as a glucose-dependent secretagogue. Genetic, genomic, and pharmacologic analyses established that the molecular clock controls the expression of the purinergic P2Y1 receptor to mediate the insulinotropic activity of ivermectin. These findings identify the P2Y1 purinergic receptor as a target to rescue circadian β-cell failure and establish a chemical genetic screen for endocrine therapeutics.

## INTRODUCTION

Type 2 diabetes is an escalating epidemic caused by interactions between genes and the environment, resulting in the co-occurrence of β-cell failure in the setting of insulin resistance. Recent epidemiologic evidence has shown that shift work and sleep disturbance are risk factors for diabetes (1), while genetic studies have revealed that disruption of the circadian clock within cells of the endocrine pancreas leads to impaired insulin exocytosis and hypoinsulinemic hyperglycemia (2, 3). At the molecular and cellular level, the circadian clock is composed of a transcriptional feedback loop in which CLOCK/BMAL1 in the forward limb drive the expression of the repressors PER1/2/3 and CRY1/2 within the negative limb that feedback to inhibit CLOCK/BMAL1 in a cycle that repeats itself every 24-hr. An additional stabilizing loop involving ROR/REV-ERB regulates BMAL1 expression (4). Recent small molecule screens have identified an expanded repertoire of factors that modulate the core clock, including casein kinase 1 inhibitors that lengthen the circadian period through regulation of the PER proteins (5, 6) and direct modulators of cryptochrome stability that exhibit glucoregulatory properties *in vivo* (7). Clock modulators may also impact metabolic homeostasis at the whole animal level (8), though achieving specificity for nuclear receptors via small molecule approaches remains challenging (9). One intriguing possibility is that small molecules may be leveraged to repair defects downstream of circadian disruption rather than through manipulation of the core clock itself.

Given that cell-intrinsic disruption of the circadian clock in mammals leads to β-cell failure (2, 10, 11), we sought to determine whether specific methods to correct insulin secretory defects downstream of the circadian clock might identify key pathways in insulin secretion and new therapeutic targets. To this end, we implemented a high-throughput luminescence-based screen of a library of 2,640 drug or drug-like compounds for the induction of insulin secretion in the context of β-cell circadian gene disruption. Our results identified the macrolide ivermectin as an insulin secretagogue which activates the P2Y1 purinergic receptor. We further identified the P2Y1 receptor as a direct transcriptional target of the molecular clock and a potent regulator of glucose-dependent calcium signaling and metabolism. Our findings establish a chemical genetic strategy to identify novel endocrine cell therapeutics.

## RESULTS

### High-throughput screen for chemical modulators of insulin secretion in circadian mutant β cells

Based upon our finding that circadian genes regulate insulin secretion and β-cell survival, we developed a phenotype-driven chemical genetic screening platform to identify small molecules that enhance insulin secretion in a cell-based model of circadian β-cell failure (**Fig 1A**). We previously generated clonal *Bmal1*^*-/-*^ Beta-TC-6 β-cell lines that eliminate an exon encoding the basic-helix-loop-helix (bHLH) DNA binding domain (11). We found that these BMAL1-ablated β-cell lines recapitulate the secretory defects observed in primary clock-deficient islets (2, 10). We next generated stable WT and *Bmal1*^*-/-*^ *β*-cell lines with a luciferase readout for insulin secretion using an insulin-NanoLuciferase (NanoLuc)-expressing lentivirus (**Fig 1B**). We validated the direct correspondence between insulin-NanoLuc bioluminescence and levels of peptide secretion under increasing physiologic concentrations of glucose (2–20 mM) (R^2^=0.8937) (**Fig 1C**). We further confirmed impaired insulin secretion by reduced bioluminescence in *Bmal1*^*-/-*^ compared to WT β-cell lines expressing insulin-NanoLuc in response to stimulatory concentrations of glucose (20 mM), potassium chloride, forskolin, and the phosphodiesterase inhibitor 3-isobutyl-1-methylxanthine (IBMX) (**Fig 1D**). We also validated the use of the diacylglycerol mimetic phorbol 12-myristate 13-acetate (PMA) as a positive control for the screen (Z’-factor score 0.69) (**Fig 1D-F**) (10).

**Figure 1.**
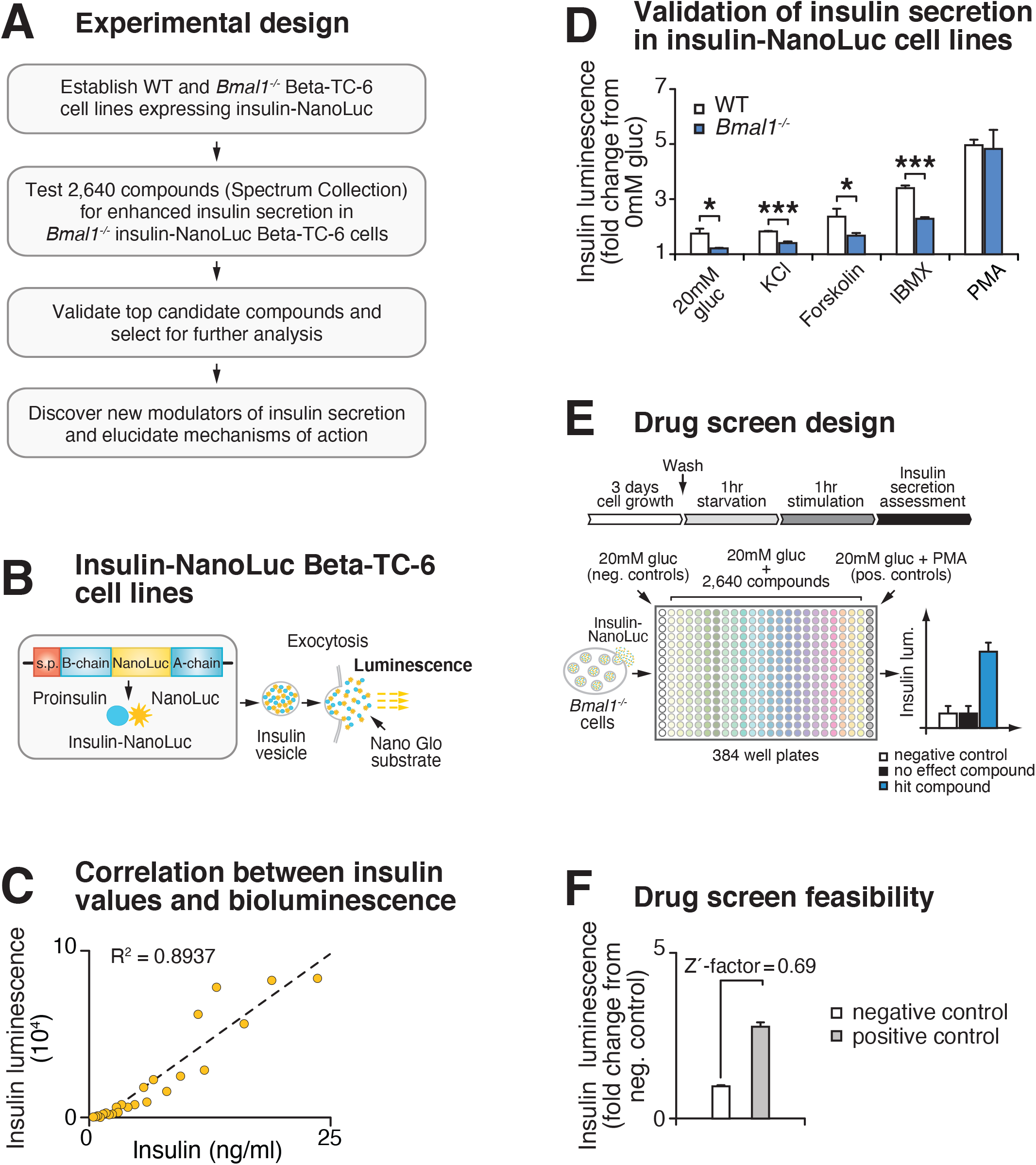
High-throughput screen for chemical modulators of insulin secretion in circadian mutant β cells. **(A)** Flow chart of “phenotype”-driven cell-based genetic screening platform to identify molecules and pathways that enhance insulin secretion during circadian β-cell failure. **(B)** Schematic of insulin-NanoLuciferase (NanoLuc) fusion construct, with bioluminescence as a proxy for insulin secretion. **(C)** Correlation between insulin-NanoLuc bioluminescence and insulin values measured by ELISA in response to a range of glucose concentrations (2-20 mM) (R^2^=0.8937). **(D)** Insulin-NanoLuc bioluminescence following 1-hr exposure to 20 mM glucose, 30 mM KCl, and 20 mM glucose plus 2.5 µM forskolin, 500 µM IBMX, or 10 µM PMA in WT and *Bmal1*^*-/-*^ insulin-NanoLuc Beta-TC-6 cells (n=3-10 repeats/condition). **(E)** Insulin-NanoLuc-expressing Beta-TC-6 *Bmal1*^*-/-*^ cells were plated in nine 384 well plates prior to exposure to 10 µM of each of the 2,640 compounds from the Spectrum collection in combination with 20 mM glucose. Negative (20 mM glucose alone) and positive (20 mM glucose plus 10 µM PMA) controls were included on each plate. **(F)** Drug screen feasibility test comparing negative (20 mM glucose only) and positive (20 mM glucose plus PMA) controls (n=3 repeats) (Z’-factor = 0.69). All values represent mean ± SEM. * p<0.05, *** p<0.001.

### Identification and validation of high throughput screen lead compounds in murine islets at high and low glucose concentrations

We next used insulin-NanoLuc-expressing *Bmal1*^*-/-*^ β-cell lines to screen 2,640 drugs and drug-like molecules from the Spectrum Collection (MicroSource Discovery Systems, Inc, New Milford, CT) to identify compounds that enhance insulin secretion (**Fig 1E**). Insulin-NanoLuc-expressing *Bmal1*^*-/-*^ Beta-TC-6 cells were plated at 40,000 cells/well in a total of nine 384-well plates, incubated for 3 days, and then treated for 1 hr with either (i) 20 mM glucose alone (negative control which elicits reduced insulin secretion in *Bmal1*^*-/-*^ cells), (ii) 20 mM glucose plus 10 µM of one of the 2,640 compounds, or (iii) 20 mM glucose plus 10 µM PMA (positive control known to enhance insulin secretion in both *Bmal1*^*-/-*^ mouse islets and Beta-TC-6 cells) (10). Luciferase intensity from the supernatant was measured following exposure to NanoGlo Luciferase Assay Substrate (**Fig 1E**).

We initially identified 19 hit compounds that both significantly enhanced insulin secretion and elicited a response of greater than 3 standard deviations from the mean (Z score > 3) with more than a 1.25-fold increase, exceeding the upper 99% confidence interval of the negative control (**Fig 2A, Fig S1A, Table S1**). Of these, seven were excluded from further analysis because of reported toxic effects or lack of availability of the compound (**Fig S1A**). The remaining 12 hit compounds mediate activity of ligand-gated cell surface receptors and ion channels that stimulate second messenger signaling cascades (**Fig 2B-C**) (12, 13). Of these, four target ion channels (tacrine hydrochloride, suloctidil, dyclonine hydrochloride, and ivermectin) (**Figs 2B-C**) (14-23). Five target seven-transmembrane G-protein coupled receptors (GPCRs) that signal through phospholipase C (PLC) and diacylglycerol (DAG) to activate insulin secretion and β-cell gene transcription (benzalkonium chloride, carbachol, isoetharine mesylate, pipamperone, and ivermectin) (**Figs 2B-C**) (17, 24-30). Similar to the hit compounds of our screen, our previous results showed that carbachol, a muscarinic G_q_-coupled receptor agonist, and the DAG mimetic PMA rescue insulin secretion in *Bmal1*^*-/-*^ islets (10). Four additional hit compounds act as acetylcholinesterase inhibitors, promoting enhanced glucose-dependent insulin secretion in response to acetylcholine through the muscarinic GPCRs, as well as the ionotropic nicotinic acetylcholine receptors (tyrothricin, tomatine, carbachol, and tacrine hydrochloride) (**Figs 2B-C**) (31-36). One compound has been shown to promote insulin secretion by inhibition of the mitochondrial protein tyrosine phosphatase PTPM1 (alexidine hydrochloride) (**Figs 2B-C**) (37, 38), and another likely affects β-cell function by signaling through the mineralocorticoid receptor (deoxycorticosterone) (**Figs 2B-C**) (39). Finally, in addition to ion channels and GPCRs, the macrolide ivermectin has also been shown to signal in micromolar concentrations though several ionotropic receptors, including purinergic, GABAergic, and glycine receptors, as well as through the farnesoid X nuclear receptor (17, 40, 41).

**Figure 2.**
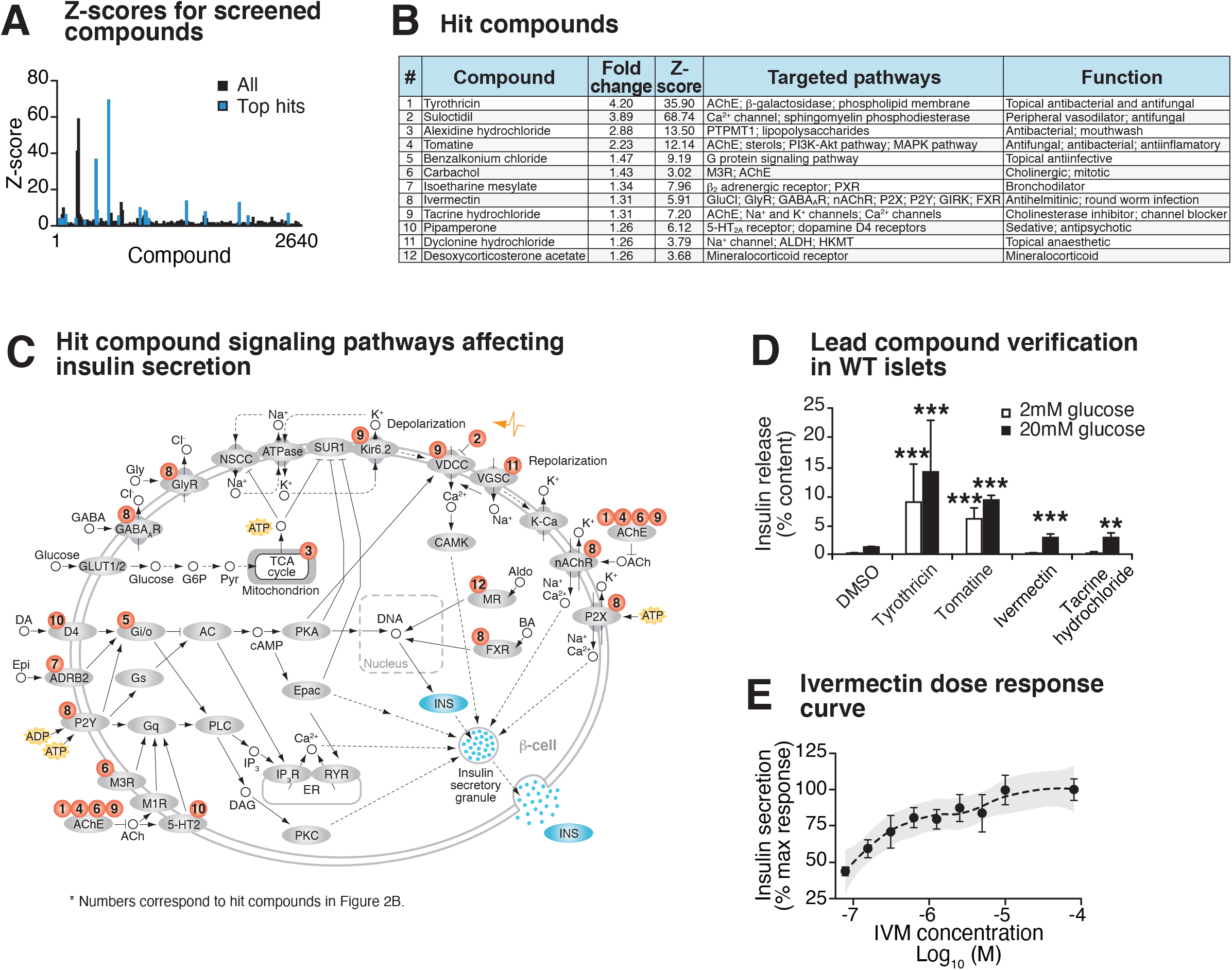
Identification and validation of high-throughput screen lead compounds in murine islets at high and low glucose concentrations. **(A)** Z-scores for all 2,640 screened compounds, with hit compounds indicated in blue. **(B)** Top 12 hit compounds identified from screen with a fold increase >1.25 and a Z-score >3 which were selected for further analysis. Known functions and published molecular pathways targeted by these compounds are indicated. **(C)** Model of potential mechanisms of action of the top 12 hit compounds to affect insulin secretion in the β cell. **(D)** Glucose-responsive insulin secretion by ELISA at 2 mM and 20 mM glucose in WT mouse islets following exposure to 4 lead candidate compounds (n=3-11 mice/compound). **(E)** IVM dose response curve (n=6-8 repeats/dose), ranging from 0.078 µM to 80 µM IVM, in insulin-NanoLuc expressing Beta-TC-6 cells. Shaded area represents 95% confidence intervals for the LOESS curve. All values represent mean ± SEM. ** p<0.01, *** p<0.001.

Ten of these twelve hit compounds were not considered for further analysis because of either the high dose required to achieve insulin secretion (**Fig S1B**) or because they augmented insulin release in low basal glucose (2 mM) in intact WT mouse primary islets (**Fig 2D**). One of the remaining compounds induces hepatotoxicity after prolonged use (tacrine hydrochloride) (42). We therefore focused our attention on ivermectin (IVM) due to its dose-dependent enhancement of glucose-stimulated insulin secretion in insulin-NanoLuc-expressing Beta-TC-6 cells, as well as its robust rescue of insulin secretion in *Bmal1*^*-/-*^ islets (**Fig 2D-E**).

### Lead compound ivermectin regulates glucose-stimulated calcium flux and insulin exocytosis in *Bmal1* mutant islets

To test whether IVM drives GSIS in β-cell lines and primary mouse islets, we first assessed the impact of both acute treatment (1-hr) and overnight exposure (24-hr) with 10 µM IVM on the ability of WT β cells and mouse islets to secrete insulin (**Figs 3A, S2A**). Consistent with our initial bioluminescence assay, we observed that IVM enhanced insulin secretion in a glucose-dependent manner following both 1-hr IVM exposure and 24-hr pre-treatment with IVM in β-cell lines and WT mouse islets, suggesting both acute and longer-term exposure to IVM enhances β-cell function (**Figs 3A, S2A**). Since there was not a significant increase in insulin secretion with overnight (∼2 fold) compared to acute (∼1.5-1.6 fold) IVM exposure, further analysis of IVM as a potentiator of insulin secretion was performed only with acute treatment.

**Figure 3.**
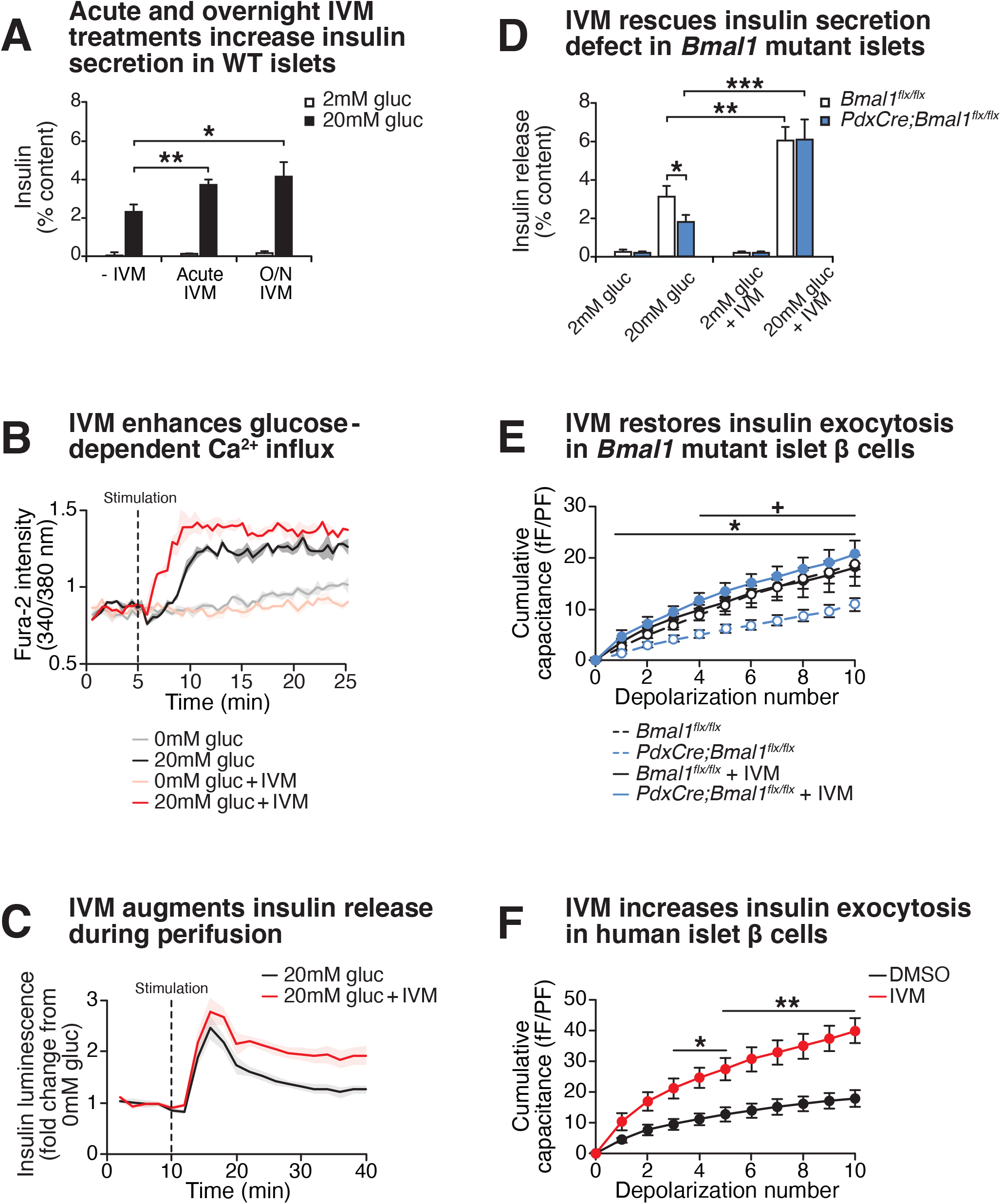
Effect of lead compound ivermectin on glucose-stimulated insulin exocytosis and calcium flux from WT and *Bmal1* mutant β cells. **(A)** Insulin secretion (expressed as % content) assessed by ELISA at 2 mM and 20 mM glucose in WT mouse islets in response to 1-hr 10 µM IVM treatment or 24-hr 10 µM IVM pre-treatment (n=5 mice). Data was analyzed by 2-way ANOVA and FDR correction for multiple testing. **(B)** Ratiometric determination of intracellular Ca^2+^ using Fura2-AM dye in WT Beta-TC-6 cells stimulated in the presence or absence of 10 µM IVM (n=3 repeats/condition). **(C)** Perifusion analysis of insulin secretion in pseudoislets from WT insulin-NanoLuc cells in response to 10 µM IVM in the presence of 20 mM glucose (n=6 repeats/condition). **(D)** Insulin secretion as assessed by ELISA from islets isolated from 8 mo old pancreas-specific *Bmal1* knockout (*Pdx-Cre;Bmal1*^*flx/flx*^) and *Bmal1*^*flx/flx*^ mice in the presence or absence of 10 µM IVM (n=10-11 mice/genotype). **(E)** Capacitance measurements in β cells from *PdxCre;Bmal1*^*flx/flx*^ and *Bmal1*^*flx/flx*^ mouse islets treated with 10 µM IVM (n=4-5 mice/genotype, 5-16 cells per mouse). Asterisks denote significance between *PdxCre;Bmal1*^*flx/flx*^ and *PdxCre;Bmal1*^*flx/flx*^ + IVM; plus symbols denote significance between *Bmal1*^*flx/flx*^ and *PdxCre;Bmal1*^*flx/flx*^ for all depolarization numbers indicated. */+ p<0.05. **(F)** Capacitance measurements in β cells from human islets treated with 10 µM IVM (n=3 donors, 7-11 cells per donor). Capacitance and calcium data were analyzed by 2-way repeated measures ANOVA with Bonferroni correction for multiple testing. All values represent mean ± SEM. * p<0.05, ** p<0.01, *** p<0.001.

Chemical energy from ATP generated by glucose metabolism within the β cell triggers closure of the sulfonylurea-linked potassium channel, depolarization of the plasma membrane, and opening of voltage-gated calcium channels leading to stimulus-secretion coupling. To assess the mechanism of IVM-induced insulin secretion, we next monitored real-time calcium influx using ratiometric fluorescence imaging in WT β cells in the presence of both glucose and IVM. We observed an immediate and robust glucose-stimulated intracellular calcium response within 2 minutes of IVM stimulation (p<0.05) (**Fig 3B**). Importantly, this effect was only observed in the presence of high glucose, consistent with results of our initial NanoLuc 384-well plate screening and subsequent ELISA-based analyses of glucose-stimulated insulin secretion. In contrast, the Ca^2+^ channel inhibitor isradipine completely suppressed Ca^2+^ influx and insulin secretion (**Fig S2D-E**) (43). To determine whether increased calcium influx corresponded with productive insulin release following IVM treatment, we used a perifusion system to directly measure NanoLuc activity in eluates harvested from IVM-treated β cells during both the first and second phase of insulin secretion (**Fig 3C**). IVM significantly increased insulin release by 12 minutes post-stimulation (p<0.05) and throughout most of the second phase of insulin secretion (>15 min), consistent with continuous release of reserve insulin granules (44) (**Fig 3C**).

Since our cell-based studies indicated that IVM stimulates GSIS within immortalized β-cell lines, we next sought to determine whether IVM restores insulin secretion in the context of circadian disruption within primary islets, which are composed of multiple hormone-releasing cell types (45). To test this idea, we administered IVM to mouse islets isolated from pancreas-specific *Bmal1*^*-/-*^ mice, revealing a 3.3-fold elevation of GSIS following exposure to the drug in the mutant islets (**Fig 3D**). To determine if IVM can improve glucose homeostasis in diabetic animals, we next tested the effects of chronic IVM administration in the well-characterized *Akita* model of β-cell failure (46). Daily intraperitoneal IVM (1.3 mg/kg body weight) was administered to *Akita* mice over a 14-day period (47), terminating in assessment of glucose tolerance and *ex vivo* GSIS. Treatment with IVM significantly improved glucose tolerance and augmented glucose-stimulated insulin release from islets isolated from these mice (**Fig S2B-C**). Given that our prior genomic and cell physiologic studies have localized the β-cell defect in circadian mutant mice to impaired insulin exocytosis (11), and as IVM augmented insulin secretion in *Bmal1* mutant islets, we next sought to determine whether IVM might enhance depolarization-induced exocytosis using electrophysiologic analyses (48). We assessed cumulative capacitance, a measure of increased cell surface area as insulin granules fuse to the plasma membrane, in β cells from islets of control and pancreas-specific *Bmal1* mutant mice, as well as from human cadaveric islets. While *Bmal1* mutant cells displayed reduced rates of exocytosis following direct depolarization (as indicated by reduced capacitance), 10 µM IVM treatment rescued the defect in *Bmal1* mutant cells, increasing cumulative capacitance from 11.0 to 20.7 fF/pF after 10 consecutive depolarization steps (**Fig 3E**). IVM treatment also enhanced cumulative capacitance in human β cells from 17.9 to 39.7 fF/pF (**Fig 3F**). Together these data show that IVM augments β-cell early calcium influx in a glucose-dependent manner to promote increased vesicle fusion and release.

### Purinergic receptor P2Y1 mediates IVM-induced insulin exocytosis

In addition to IVM, several of the predicted targets of the insulinotropic compounds from our screen involve second-messenger signaling, raising the possibility that circadian disruption may be overcome by augmenting hormonal or metabolic factors that promote peptide exocytosis. IVM is a readily-absorbable and potent derivative of avermectin B_1_ that acts to allosterically regulate several different types of cell surface receptors, including the purinergic and GABA receptors, as well as nuclear transcription factors such as the farnesoid X receptor (FXR) (47, 49-51). Since IVM augments insulin secretion in *Bmal1*^*-/-*^ cells, we hypothesized that the expression of putative IVM targets may be reduced during circadian disruption. First, through RNA-sequencing we observed significantly higher levels of expression of the transcript encoding the purinergic receptor P2Y1 (*P2ry1*) in WT β cells compared to transcripts encoding FXR or GABA components (**Fig S3A**). We further observed enrichment of BMAL1 chromatin binding within enhancer regions 266 - 41 kb upstream of the *P2ry1* gene transcription start site by chromatin immunoprecipitation-sequencing (GSE69889) (**Fig 4A**), as well as rhythmic expression of *P2ry1* in wild-type Beta-TC-6 pseudoislets (**Fig S3B**). Finally, we observed a 3.1-fold reduction (Adj. P = 10^−55^) in expression of *P2ry1* in circadian mutant β cells (**Figs 4A, S3A**) (GSE146916), suggesting a direct role of the circadian clock in *P2ry1* expression.

**Figure 4.**
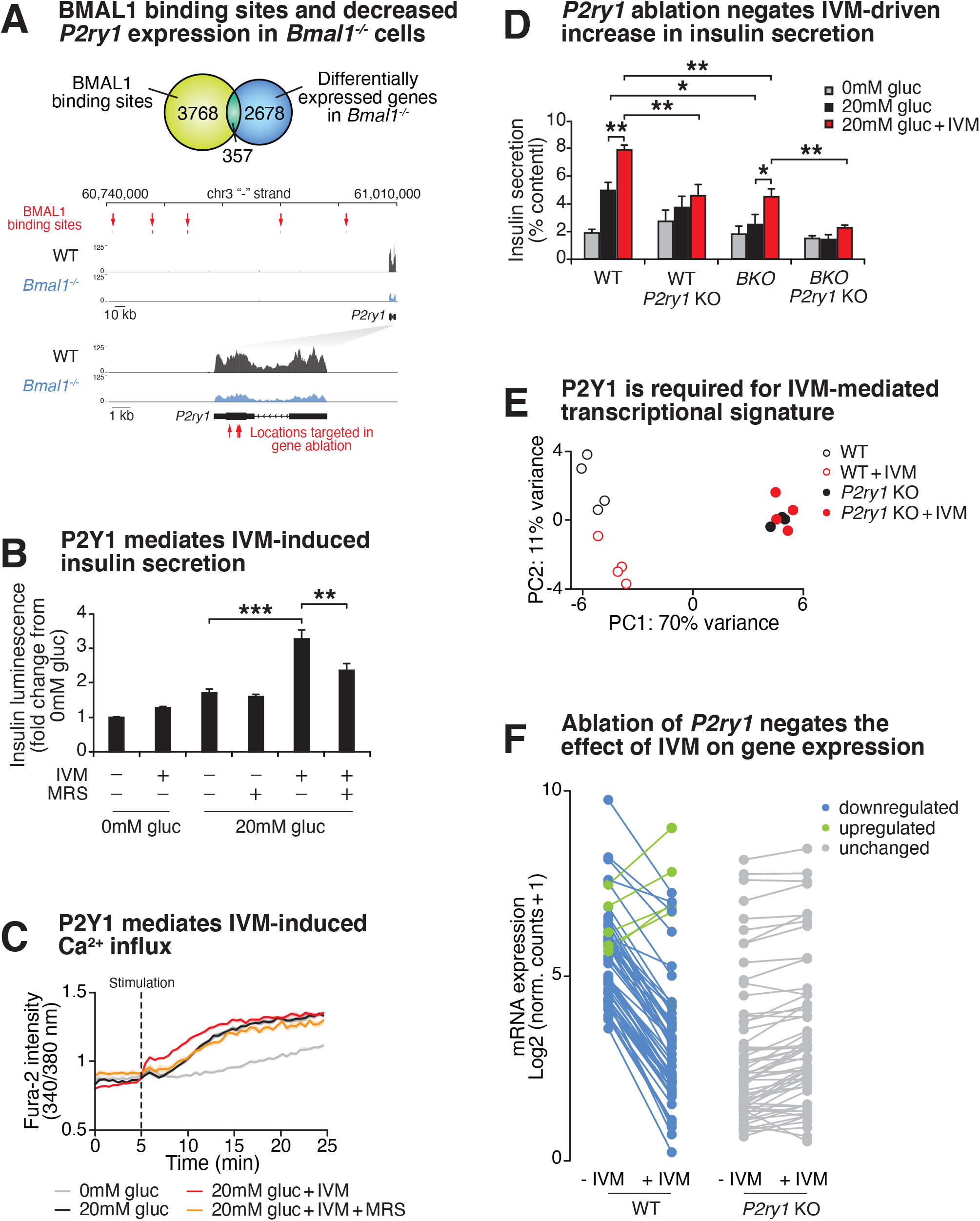
Purinergic receptor P2Y1 is required for IVM to augment insulin exocytosis. **(A)** Venn diagram of BMAL1 binding sites identified by ChIP-sequencing overlapping with differentially-expressed genes identified by RNA-sequencing in *Bmal1*^*-/-*^ *β*-cell line compared to control cell line (*top*). Browser tracks showing decreased expression of *P2ry1* gene in *Bmal1*^*-/-*^ cells compared to controls. BMAL1 binding sites upstream of the *P2ry1* gene are also indicated (*bottom*). **(B)** Bioluminescence from WT insulin-NanoLuc pseudoislets in response to 10 µM IVM and/or 10 µM of the P2Y1 antagonist MRS2179 (n=4-8 experiments, 3-8 repeats per experiment). **(C)** Ratiometric determination of intracellular Ca^2+^ using Fura2-AM dye in WT Beta-TC-6 cells stimulated in the presence or absence of 10µM IVM (n=3-8 experiments, 4-12 repeats per experiment). **(D)** Insulin secretion by ELISA in pseudoislets from *P2ry1* KOs and control WT and *Bmal1*^*-/-*^ Beta-TC-6 cells (n=4/genotype/condition). Benjamini and Hochberg FDR-adjusted P values were computed for multiple comparisons following two-way ANOVA. **(E)** First two principal components (PC1 and PC2) following unbiased principal component analysis (PCA) of DESeq2 normalized counts in WT, WT + IVM, *P2yr1* KO, and *P2yr1* KO cells (n=4 per group). **(F)** Mean log_2_-transformed DESeq2-normalized counts in WT, WT + IVM, *P2yr1* KO, and *P2yr1* KO cells (n=4 per group) at differentially-expressed (1.5 fold, adjusted P value < 0.05) transcripts identified between WT and WT + IVM treated cells). All values represent mean ± SEM. * p<0.05, ** p<0.01, *** p<0.001.

Based upon evidence that IVM targets purinergic receptors (52, 53), that *P2ry1* is within the top 12% of expressed transcripts in the murine β cell (by transcripts per million), and that BMAL1 specifically controls *P2ry1* amongst the purinergic receptor family in the β cell (**Figs 4A, S3A-B**), we sought to test the functional role of the P2Y1 receptor in the insulinotropic action of IVM. Pharmacologic inhibition of P2Y1 using the subtype-specific inhibitor MRS2179 in the presence of both high glucose and 10 µM IVM resulted in a 52% reduction in insulin secretion by bioluminescence and a reduction in calcium influx to levels similar to those observed during high glucose alone, as assessed by Fura2-AM ratiometric determination of intracellular calcium (**Figs 4B-C**). In addition to evidence that pharmacological blockade of P2Y1 receptor signaling abrogates IVM activity, we also tested the requirement of P2Y1 receptor signaling following CRISPR-Cas9-mediated knockout of the P2Y1 receptor in both WT and *Bmal1*^*-/-*^ *β* cells (**Fig S4A**). While IVM enhanced glucose-stimulated insulin secretion in WT and *Bmal1*^*-/-*^ *β* cells by 60% and 80%, respectively, IVM did not significantly enhance glucose-stimulated insulin secretion in cells lacking the P2Y1 receptor (**Fig 4D**). Similar to the pharmacologic findings with the P2Y1 antagonist MRS2179, these results demonstrate a requirement for P2Y1 in IVM-induced GSIS. P2Y1R signaling involves activation of Ca^2+^ entry and intracellular release, which results in both acute stimulation of insulin granule trafficking and activation of transcription factors that may be involved in β-cell function (54-56). To analyze gene expression changes induced by P2Y1 activation, we performed RNA-sequencing to compare the IVM response within both WT and *P2ry1*^*-/-*^ β cells following stimulation with glucose or glucose plus IVM. Principal component analysis (PCA) was performed using log-transformed count data from the top 500 most variable genes across all samples (57). This revealed distinct patterns in mRNA expression between IVM- and control-treated WT cells along PC2, while there was no separation between IVM- and control-treated *P2ry1*^*-/-*^ β cells, suggesting that P2Y1 is required for IVM-mediated transcriptional changes in β cells (**Fig 4E**). In WT cells, IVM induces differential expression of 65 transcripts (1.5-fold change, Adj. P value < 0.05), including up-regulation of the immediate early gene *Fos* (58) and down-regulation of *Aldolase B*, whose expression has been linked to reduced insulin secretion in human islets (59) (**Figs 4F, S4B**). Strikingly, none of these transcripts were significantly altered by IVM in the *P2ry1*^*-/-*^ β cells (all adjusted P value > 0.05) (**Figs 4F, S4B**). Taken together, these data suggest that the circadian clock program controls P2Y1 expression to modulate glucose-stimulated insulin secretion and highlight the utility of a genetic-sensitized drug screen for identification of therapeutic targets in circadian dysregulation and diabetes.

## DISCUSSION

We have identified an unexpected role for the P2Y1 receptor as a BMAL1-controlled insulinotropic factor required for enhanced β-cell glucose-stimulated Ca^2+^ influx and insulin secretion in response to IVM. While P2Y receptors have been previously implicated in calcium and insulin secretory dynamics in β cells, modulation has been primarily demonstrated using agonists that mimic ATP/ADP derivatives that have deleterious effects on thrombosis (54-56, 60). Little is known about P2Y1 targeting in disease states, such as circadian disruption and/or type 2 diabetes, or whether P2Y1 is controlled at a transcriptional level. Our evidence that P2Y1 is expressed under control of the circadian clock derives from analyses at the level of both chromatin binding by the core clock factor BMAL1 and genome-wide differential RNA expression analysis in circadian mutants. Intriguingly, P2X and P2Y receptors are required for Ca^2+^ signaling in the suprachiasmatic nucleus (61, 62), yet their role in circadian regulation of peripheral tissues has not been well studied. Our pharmacological and genetic analyses are the first to reveal that enhancement of P2Y1 receptor activity can bypass the transcriptional deficits exhibited in circadian mutant β cells and restore insulin secretion. Future studies will be required to determine the precise mechanism by which IVM modulates P2Y1 activity. One possibility is that IVM may augment P2X-P2Y1 crosstalk to drive insulin secretion, which has been shown to drive Ca^2+^ and P2Y1-dependent activation of other cell types (52, 63).

Previous physiologic and transcriptomic studies have shown that circadian regulation of insulin exocytosis involves control of the expression and activity of cell-surface receptors and second messenger systems (10, 64). We based our drug screen on the idea that modulators of insulin secretion in cells that lack a functional clock would complement prior genomic analyses revealing circadian control of peptidergic hormone exocytosis and also to provide proof-of-principle that the clock can be leveraged to sensitize screening for new chemical modulators of β-cell function. This approach identified Ca^2+^-dependent pathways as a potential route to ameliorate circadian disruption and to enhance glucose-stimulated insulin secretion. Importantly, several of the compounds identified in our screen have been used in disease treatment and have known mechanisms of action, including the cholinergic activators carbachol and tacrine (65, 66). The identification of these compounds in our screen raises the intriguing possibility of using drug derivatives related to these molecules for type 2 diabetes treatment, particularly in the context of circadian/sleep disruption.

The study of transcriptional rhythms across the 24-hr circadian cycle has previously revealed a diverse landscape of clock-controlled genes and pathways (67). Despite the identification of thousands of tissue-specific and clock-controlled transcripts, limited advances have been made in utilizing this information to treat diseases associated with circadian disruption, including type 2 diabetes. One approach to this challenge has been to intervene and restore the molecular clock program using pharmacology (Nobiletin) (8), micronutrient supplementation (NAD^+^ precursors) (68, 69), or enforced behavioral rhythms (such as time restricted feeding) (70). However, it remains unclear how altering the whole-body clock will affect nutritional and hormonal dynamics at a cellular level. Another approach has been to directly target clock-controlled genes with known function in health and disease (71), or to look at gain/loss of circadian control in health versus disease (72). This approach requires an understanding of gene function within a given tissue, and thus limits the identification of novel therapeutic targets. In the studies performed here we sought to address the challenge of connecting clock control of transcription with druggable targets by using an unbiased small molecule drug screen, in tandem with functional genomics, to elucidate mechanisms of insulin secretory dynamics. Since the circadian timing system has been shown to not only regulate the function of mature β cells, but also the regenerative capacity of islets in both the context of the mouse (73) and in human embryonic stem cell differentiation (74), molecules identified in cell-based genetic screens may provide broad applicability as therapeutics.

## Materials and Methods

### Reagents

Ivermectin, (+)-Bicuculline, and MRS2179 tetrasodium salt were obtained from Tocris (R&D Systems, Inc, Minneapolis, MN). Isradipine was purchased from Cayman Chemical Company (Ann Arbor, MI). Exendin-4, PMA, guggulsterone, carbamoylcholine chloride (carbachol), forskolin, tyrothricin, alexidine hydrochloride, and benzalkonium chloride were obtained from Sigma-Aldrich (St. Louis, MO). Suloctidil, tomatine, isoetharine mesylate, tacrine hydrochloride, pipamperone, dyclonine hydrochloride, and desoxycorticosterone acetate were purchased from MicroSource Discovery Systems.

### Animals

Male WT C57BL6J mice and C57BL/6-*Ins2*^*Akita*^/J mice were purchased from the Jackson Laboratory (Bar Harbor, ME). *PdxCre;Bmal1*^*flx/flx*^ mice were produced and maintained on C57BL6J background at Northwestern University Center for Comparative Medicine (75). Unless otherwise stated, animals were maintained on a 12:12 light:dark cycle and allowed free access to water and regular chow. All animal care and use procedures were conducted in accordance with regulation of the Institutional Animal Care and Use Committee at Northwestern University.

### Cell Culture

Beta-TC-6 cells were obtained from ATCC (Manassas, VA) (CRL-11506), and *Bmal1*^*-/-*^ Beta TC-6 β-cell lines were previously derived as described (11). Cells were cultured in Dulbecco’s Modified Eagle’s Medium (DMEM; Gibco, Aramillo, TX) supplemented with 15% fetal bovine serum (BioTechne, Minneapolis, MN), 1% penicillin-streptomycin (Gibco), and 1% L-glutamine (Gibco) at 37°C with 5% CO_2_. Culture medium was exchanged every 2-3 days. All cells used in experiments were at <15 passages.

### Generation of WT and *Bmal1*^*-/-*^ Beta-TC-6 cells stably expressing insulin-NanoLuc

We used the proinsulin-NanoLuc plasmid (David Altshuler, Addgene plasmid #62057) to provide a low cost, scalable, and rapid method to detect insulin secretion. The gene encoding NanoLuciferase was cloned into the C-peptide portion of mouse proinsulin such that cleavage within insulin vesicles by pH-sensitive prohormone convertase results in the co-secretion of NanoLuc with endogenous insulin in a stimulus-dependent manner (76). The pLX304 lentivirus packaging plasmid containing the proinsulin-NanoLuc construct was transfected into HEK293T (ATCC CRL-11268) cells with pCMV-VSVG (envelope vector) and 8.91 (packaging vector) (obtained from Jeff Milbrandt, Washington University in St. Louis). Supernatant containing lentivirus particles was harvested 48 hrs after transfection. Beta-TC-6 and *Bmal1*^*-/-*^ Beta TC-6 cells were infected with insulin-NanoLuc lentivirus, and stably expressing cells were selected by treating with puromycin (2 µg/ml, 2 days).

### CRISPR-mediated *P2ry1* deletion in WT and *Bmal1*^*-/-*^ Beta TC-6 cells

Exon 1 of the mouse *P2yr1* gene was deleted in WT and *Bmal1*^*-/-*^ Beta-TC-6 cells by CRISPR-Cas9 and homology-directed repair (HDR). Cells were co-transfected with guide RNA, P2Y1 CRISPR/Cas9 KO, and P2Y1 HDR plasmids (Santa Cruz Biotechnology, Dallas, TX) by Lipofectamine 2000 (Thermo Fisher Scientific, Amarillo, TX). After 48 hrs of transfection, stably-integrated clones were selected for puromycin resistance (puromycin dihydrochloride, Sigma-Aldrich). RNA and protein were extracted from these colonies and *P2ry1* expression was assessed by qPCR and Western blot.

### High-throughput screen for drugs to restore insulin secretion in *Bmal1*^*-/-*^ β cells and insulin secretion assays

The Spectrum Collection small molecule compound library (MicroSource Discovery Systems, Inc), which consists of 2,640 known drugs and drug-like molecules, was screened for compounds that augment insulin secretion in *Bmal1*^*-/-*^ Beta-TC-6 cells. Insulin-NanoLuc-expressing *Bmal1*^*-/-*^ Beta-TC-6 cells (40,000 cells/well) were placed into 384 well plates and cultured for 3 days at 37°C and 5% CO_2_. The cells were washed once and incubated in KRB buffer containing 0mM glucose for 1 hr. Then, KRB buffer containing 20 mM glucose in addition to the small molecules (10 µM) were added, and the cells were incubated for 1 hr. As a negative control, 16 wells received KRB buffer with only 20 mM glucose, which fails to elicit appropriate insulin secretion in *Bmal1*^*-/-*^ cells, and as a positive control, 16 wells received KRB buffer containing 20 mM glucose and 10µM PMA, which is known to induce insulin secretion in both *Bmal1*^*-/-*^ mouse islets and Beta-TC-6 cells (10). After 1 hr, the supernatant was collected and centrifuged at 500g for 30 min. The supernatant was transferred into a fresh 384-well assay plate containing NanoGlo Luciferase Assay Substrate (Promega, Madison, WI), and luciferase intensity was measured by EnSpire Plate Reader (PerkinElmer, Waltham, MA) within 30 minutes. All liquids for the high-throughput screen were dispensed using Tecan Fluent Automated Liquid Handling Platform (Tecan, Mannedorf, Switzerland) at the High-Throughput Analysis Laboratory at Northwestern University. Screen feasibility was determined by calculating Z’-factor using the following formula: Z’-factor = 1-3(σ_p_ + σ_n_) / (µ_p_ - µ_n_) (where σ_p_ is the standard deviation of positive control, σ_n_ is the standard deviation of negative control, µ_p_ is the mean intensity of positive control, and µ_n_ is the mean intensity of the negative control).

### Determination of hit compounds

Z scores for luciferase intensities produced by screened compounds were calculated from the following formula: z = (X - µ) / σ (where z is the Z score, X is the intensity of the compounds, µ is the intensity of negative control (20mM glucose), and σ is the standard deviation of negative control). A row-based correction factor was applied to all luciferase readings to adjust for logarithmic signal decay. Hit compounds were defined as those that elicited a response of greater than 3 standard deviations from the mean (Z score > 3) and more than 1.25-fold increase compared to negative control, which is the cut-off for ∼10% chance of the observation occurring by random chance. Validated hit compounds that augmented insulin secretion at low drug dose were considered lead compounds.

### Insulin secretion assays in pancreatic islets, pseudoislets, and cell lines

Mouse pancreatic islets were isolated via bile duct collagenase digestion (*Collagenase P*, Sigma) and Biocoll (Millipore) gradient separation and left to recover overnight at 37°C in RPMI 1640 with 10% FBS, 1% L-glutamine, and 1% penicillin/streptomycin. For insulin release assays, duplicates of 5 equally-sized islets per mouse were statically incubated in Krebs-Ringer Buffer (KRB) at 2 mM glucose for 1 hr and then stimulated for 1 hr at 37°C with 2 mM or 20 mM glucose in the presence or absence of 10 µM of each compound. Supernatant was collected and assayed for insulin content by ELISA (Crystal Chem Inc, Elk Grove Village, IL). Islets were then sonicated in acid-ethanol solution and solubilized overnight at 4°C before assaying total insulin content by ELISA. For insulin release assays from pseudoislets, 3 × 10^6^ cells were plated for 3 days in 60 mm suspension dishes and allowed to form pseudoislets for 2-3 days. Glucose-responsive insulin secretion was performed as described above, using 10 pseudoislets per sample and a basal glucose level of 0 mM glucose instead of 2 mM. For secretion from insulin-NanoLuc cell lines, 1 × 10^5^ cells were cultured on poly-L-lysine coated 96 well plates for 2-3 days, starved for 1 hr in 0 mM glucose KRB, then stimulated with indicated compounds and/or receptor antagonists for 1 hr in conjunction with indicated glucose concentrations. Luciferase intensity after addition of NanoGlo to supernatant was measured by Cytation3 Plate Reader (BioTek, Winooski, VT).

### Perifusion of pseudoislets

Perifusion of 100 insulin-NanoLuc pseudoislets was performed using a Biorep Technologies Perifusion System Model PERI-4.2 with at a rate of 100 µL/min KRB (0.2% BSA). After 1 hour of preincubation and equilibration at a rate of 100 µL/min with 0 mM KRB, 0 mM glucose KRB was perifused for 10 minutes, followed by perifusion for 30 minutes with 20 mM or 20 mM plus IVM. Perifusate was collected in 96 well plates and analyzed for NanoLuc activity using NanoGlo Luciferase Assay Substrate (Promega) per manual instructions, substituting lysis buffer for KRB perifusate.

### *In vivo* ivermectin treatment and glucose measurements

Mice were injected intraperitoneally for 14 days with 1.3 mg/kg body weight of IVM, which was dissolved in 40% w/v 2-hydroxypropyl-β-cyclodextrin (Sigma-Aldrich) (47). At the end of IVM treatment, mice were fasted for 14 hrs and glucose tolerance tests were performed at ZT2 following intraperitoneal glucose injection at 2g/kg body weight. Plasma glucose levels were measured by enzymatic assay (Autokit Glucose, Wako-Fujifilm, Cincinnati, OH).

### Synchronization, RNA isolation, and qPCR mRNA quantification

Where indicated, circadian synchronization was performed using 200 WT pseudoislets by first exposing cells to 10 µM forskolin for one hour, followed by transfer to normal media and RNA collection every 4 hrs 24-44 hrs following forskolin synchronization pulse. RNA was extracted from Beta-TC-6 cells and pseudoislets using Tri Reagent (Molecular Research Center, Inc, Cincinnati, OH) and frozen at ™80°C. RNA was purified according to the manufacturer’s protocol using the Direct-zol™ RNA Microprep kit (Zymo Research, Irvine, CA) with DNase digestion. cDNAs were then synthesized using the High Capacity cDNA Reverse Transcription Kit (Applied Biosystems, Amarillo, TX). Quantitative real-time PCR analysis was performed with SYBR Green Master Mix (Applied Biosystems) and analyzed using a Touch™ CFX384 Real-Time PCR Detection System (Bio-Rad, Hercules, CA). Target gene expression levels were normalized to *β-actin* and set relative to control conditions using the comparative C_T_ method. Primer sequences for qPCR as follows: *β-actin* Forward: 5′-TGCTCTGGCTCCTAGCACCATGAAGATCAA-3′, Reverse: 5′-AAACGCAGCTCAGTAACAGTCCGCCTAGAA-3; *P2ry1* Forward: 5′-TTATGTCAGCGTGCTGGTGT -3′, Reverse: 5′-ACGTGGTGTCATAGCAGGTG -3.

### RNA-sequencing and analysis

Following RNA isolation (described above), RNA quality was assessed using a Bioanalyzer (Agilent, Santa Clara, CA), and sequencing libraries were constructed using a NEBNext® Ultra™ Directional RNA Library Prep Kit for Illumina (New England BioLabs, Ipswich, MA, E7420L) according to the manufacturer’s instructions. Libraries were quantified using a NEBNext® Library Quant Kit for Illumina (New England BioLabs, E7630L) and sequenced on either an Illumina NextSeq 500 instrument using 42bp paired-end reads. For differential expression analysis, RNA raw sequence reads were aligned to the reference genome (mm10) using STAR version 2.7.2b, and raw and transcripts per million (TPM) count values determined using RSEM version 1.3.3. Differentially expressed RNAs were identified by an FDR-adjusted P value <0.05 and a fold change > 1.5 using DESeq2 version 1.32.0 in R 4.1.0. Heatmaps were generated using the pheatmap package in R. Raw mRNA sequencing data and gene abundance measurements have been deposited in the Gene Expression Omnibus under accession GSE186469.

### Intracellular calcium determination

Beta-TC-6 cells were plated at a density of 100,000 cells per well in black 96-well plates with clear bottoms and cultured overnight at 37°C and 5% CO_2_. Cells were then washed with BSA-free KRB buffer with no glucose and loaded with 5 µM Fura-2 (Invitrogen, Amarillo, TX) and 0.04% Pluronic F-127 (Invitrogen) for 30 min at 37°C. Following a wash with BSA-free KRB, Fura-2 intensity was measured after stimulation with either glucose alone or glucose plus the indicated compounds. Cells were alternately excited with 340 nm and 380 nm wavelength light, and the emitted light was detected at 510 nm using a Cytation 3 Cell Imaging Multi-Mode Reader (BioTek) at sequential 30-second intervals. Raw fluorescence data were exported to Microsoft Excel and expressed as the 340/380 ratio for each well.

### Patch-clamp electrophysiology

Patch-clamp measurement of exocytic responses in mouse β cells was performed as previously described (11). Human islets isolations approved by the Human Research Ethics Board (Pro00013094; Pro00001754) were performed at the Alberta Diabetes Institute Islet-Core according to methods deposited in the protocols.io repository (77). A total of three non-diabetic (ND) donors were examined in this study. Full details of donor information, organ processing, and quality control information can be assessed with donor number (donors R224, R225, and R226 in this study) at www.isletcore.ca. Dispersed human islets were cultured in low glucose (5.5 mM) DMEM media (supplemented with L-glutamine, 110 mg/l sodium pyruvate, 10% FBS, and 100 U/ml penicillin/streptomycin) in 35-mm culture dishes overnight. On the day of patch-clamp measurements, human or mouse islet cells were preincubated in extracellular solution at 1 mM glucose for 1 hr and capacitance was measured at 10 mM glucose with DMSO or 10 µM ivermectin as previously described (11). Mouse β cells were identified by cell size and by half-maximal inactivation of Na^+^ currents near -90 mV and human β cells were identified by immunostaining for positive insulin, following the experiment as described (48). Data analysis was performed using GraphPad Prism (v8.0c). Comparison of multiple groups was done by one- or two-way ANOVA followed by Bonferroni or Tukey post test. Data are expressed as means ± SEM, where P < 0.05 is considered significant.

### Western blotting

Beta-TC-6 cells lysates were isolated by treating cell pellets with RIPA buffer (Sigma-Aldrich) supplemented with 1x protease and 1x phosphatase inhibitors (Roche, Basel, Switzerland). Protein levels were quantified using DC Protein Assay (Bio-Rad), protein extracts were subject to SDS-PAGE gel electrophoresis and transferred to nitrocellulose membranes (GE Healthcare, Chicago, IL). Primary antibodies used were anti-P2Y1 (Santa Cruz, sc-377324) and anti-β-ACTIN (Cell Signaling, Danvers, MA, CST 4970).

### Statistical analysis

Results were expressed as mean ± SEM unless otherwise noted. Information on sample size, genotype, and p values is provided within each figure and figure legend. Statistical significance of capacitance, Fura2, and perifusion data was performed using a two-way analysis of variance (ANOVA) or mixed effects model (for datasets with missing values) with repeated measures followed by multiple comparison tests using a Bonferroni P value adjustment via Prism (v9.2.0). Statistical analysis was performed by unpaired two-tailed Student’s *t*-test unless otherwise indicated. P<0.05 was considered to be statistically significant. JTK_Cycle (v3) was used to determine rhythmicity in qPCR data, using a period length of 24 hours and considering a Benjamini-Hochberg (BH)-adjusted P value < 0.05 as statistically rhythmic (78).

## Acknowledgements

We thank all members of the Bass laboratory, Dr. Grant Barish, Dr. Lisa Beutler, and Dr. Richard Miller for helpful discussions and comments on the manuscript. We also thank Shun Kobayashi for technical assistance. We thank the Human Organ Procurement and Exchange (HOPE) program and Trillium Gift of Life Network (TGLN) for their work in procuring human donor pancreas for research. Finally, we especially thank the organ donors and their families for their kind gift in support of diabetes research.

## Author Contributions

B.M., B.J.W., A.T., and J.B. designed research. B.M., B.J.W., A.T., M.P., M.V.N., Y.K., C.O., J.E.M.F., and H.L. performed research. B.M., B.J.W., A.T., M.P. analyzed data. P.E.M. contributed conceptually. K.M.R., B.M., B.J.W., and J.B wrote the paper.

## Data Sharing Plans

Data in this study is publicly available in the GEO repository GSE186469.

---

**Figure S1.**
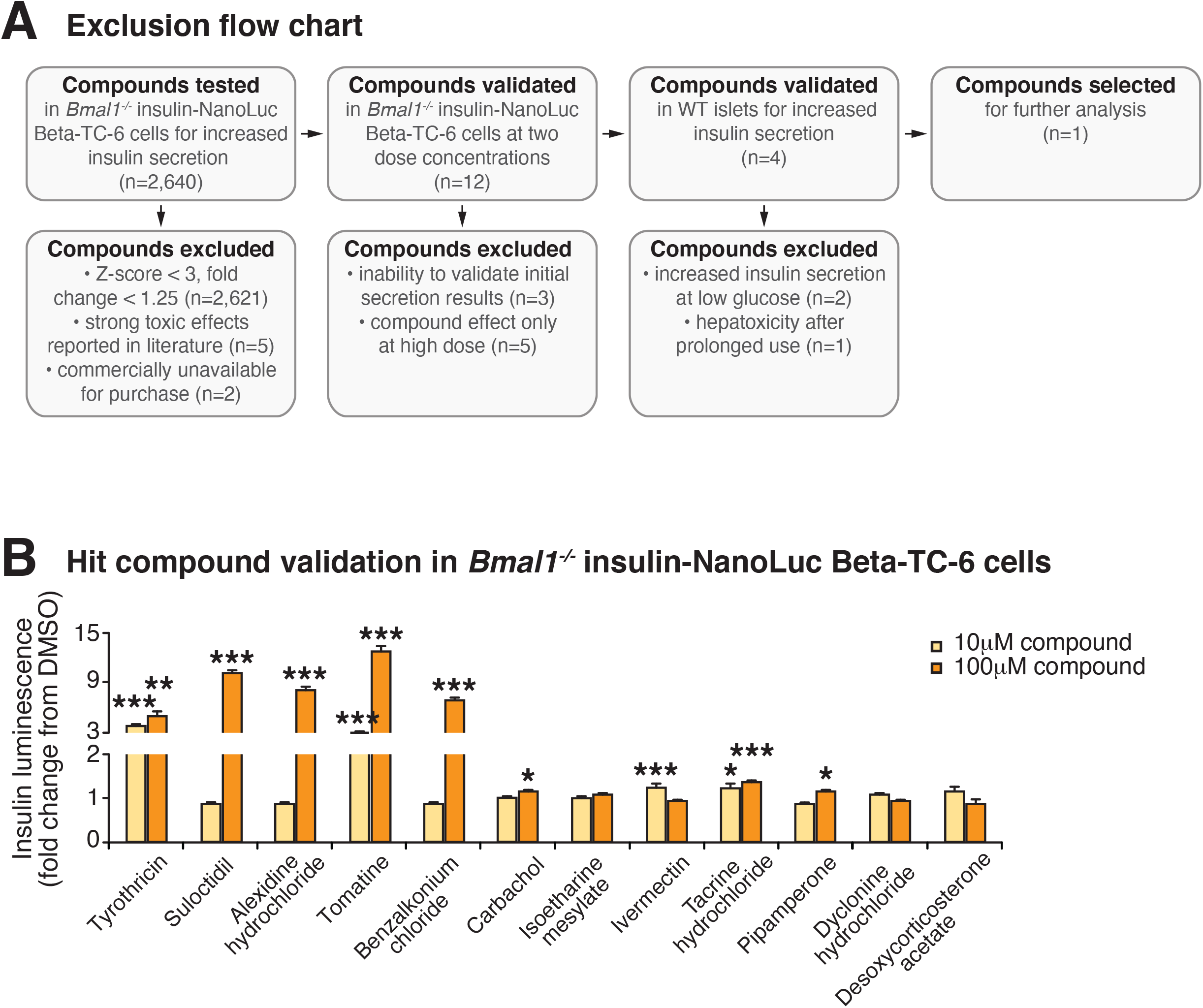
High-throughput screen for modulators of insulin secretion in circadian mutant β cells and validation of lead compounds. **(A)** Compound exclusion flow chart delineating exclusion criteria and numbers of compounds excluded at each validation step. **(B)** Hit compound validation at concentrations of 10 and 100 µM in *Bmal1*^*-/-*^ insulin-NanoLuc cells (n=3/compound). All values represent mean ± SEM. * p<0.05, ** p<0.01, *** p<0.001.

**Figure S2.**
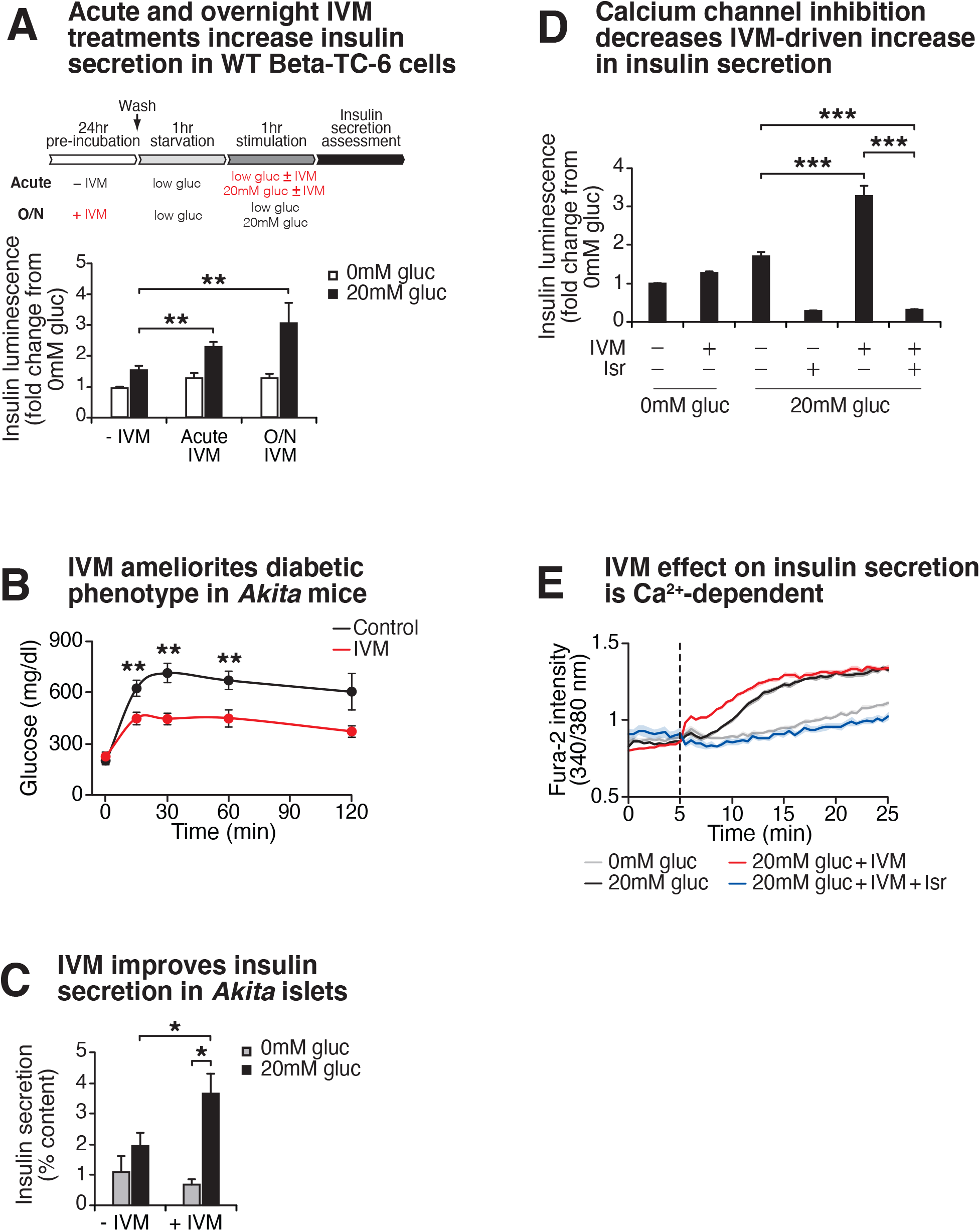
Ivermectin improves insulin exocytosis in diabetic mice. **(A)** Insulin-NanoLuc bioluminescence at 0 mM and 20 mM glucose in WT Beta-TC-6 cells in response to 1-hr 10 µM IVM treatment or 24-hr 10 µM IVM pre-treatment (n=5 experiments, 3-24 repeats/experiment). Data was analyzed by 2-way ANOVA and FDR correction for multiple testing. **(B)** Glucose levels at the indicated time points following an intraperitoneal injection of glucose (2 g/kg body weight) at ZT2 after 14 days of daily intraperitoneal injections with 1.3 mg/kg body weight of IVM (n=8 mice/genotype). Glucose levels were analyzed by 2-way repeated measures ANOVA with Bonferroni multiple testing. **(C)** Insulin secretion as assessed by ELISA from islets isolated from 4 mo old *Akita* mice in the presence or absence of 10 µM IVM (n=5-6 mice/genotype). **(D)** Insulin secretion in pseudoislets from WT insulin-NanoLuc-expressing Beta-TC-6 cells in response to 10 µM IVM and 5 µM isradipine (isr) (n=3-8 experiments, 3-8 repeats per experiment). **(E)** Ratiometric determination of intracellular Ca^2+^ using Fura2-AM dye in WT Beta-TC-6 cells stimulated with IVM or isr (n=3-8 experiments, 6-8 repeats per experiment). All values represent mean ± SEM. ** p<0.01, *** p<0.001.

**Figure S3.**
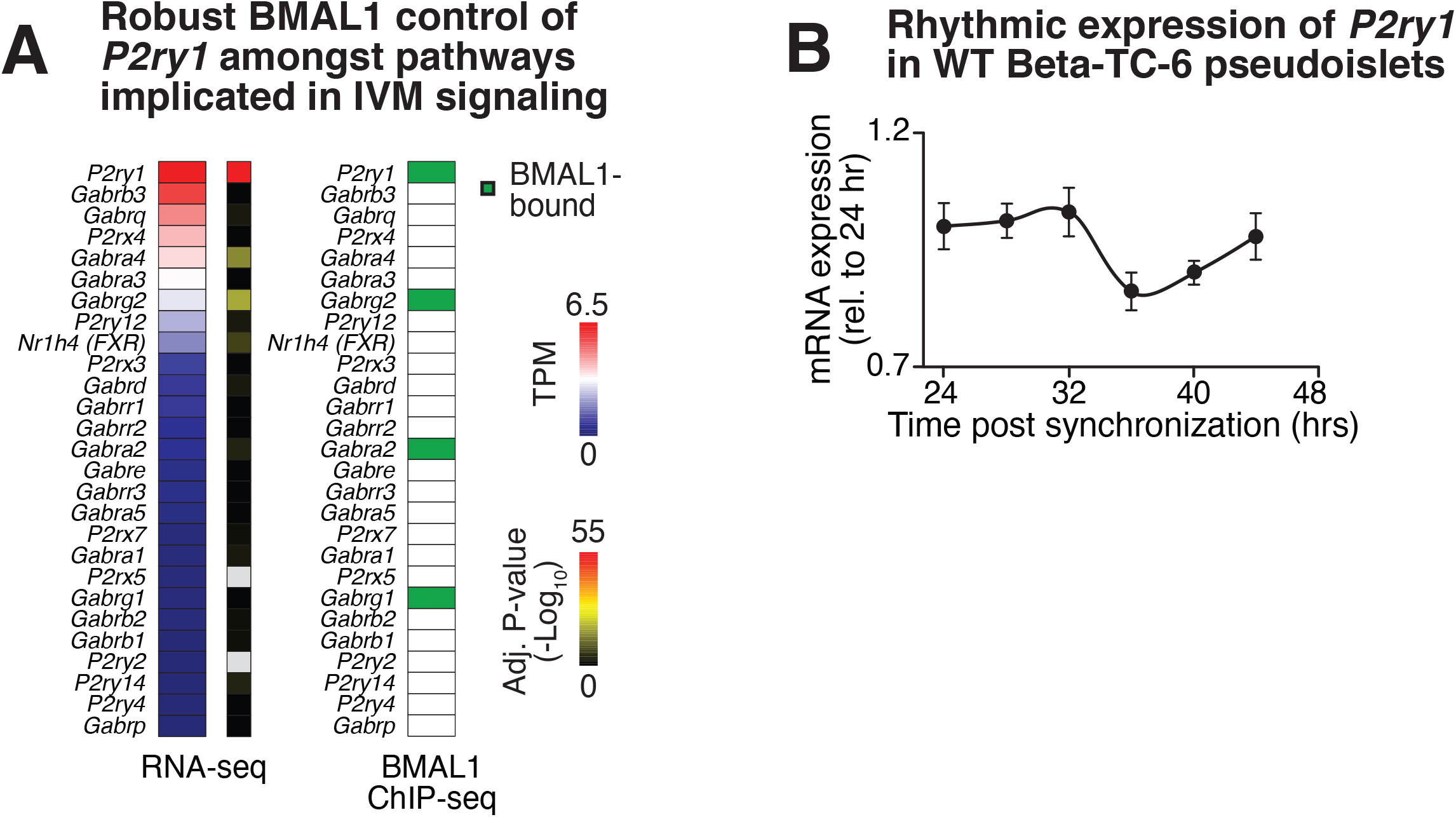
Evidence for circadian control of *P2ry1*. **(A)** mRNA abundance (TPM, transcripts per million) in WT β cells (left), DESeq2-adjusted P values from differential expression analysis in WT versus *Bmal1*^*-/-*^ *β* cells (middle), and presence or absence of an annotated BMAL1 binding site near genes of putative IVM targets (right). **(B)** Rhythmic expression of *P2ry1* gene in synchronized pseudoislets from WT Beta-TC-6 cells as assessed by quantitative real-time PCR (n=3) (FDR adjusted P value < 0.05).

**Figure S4.**
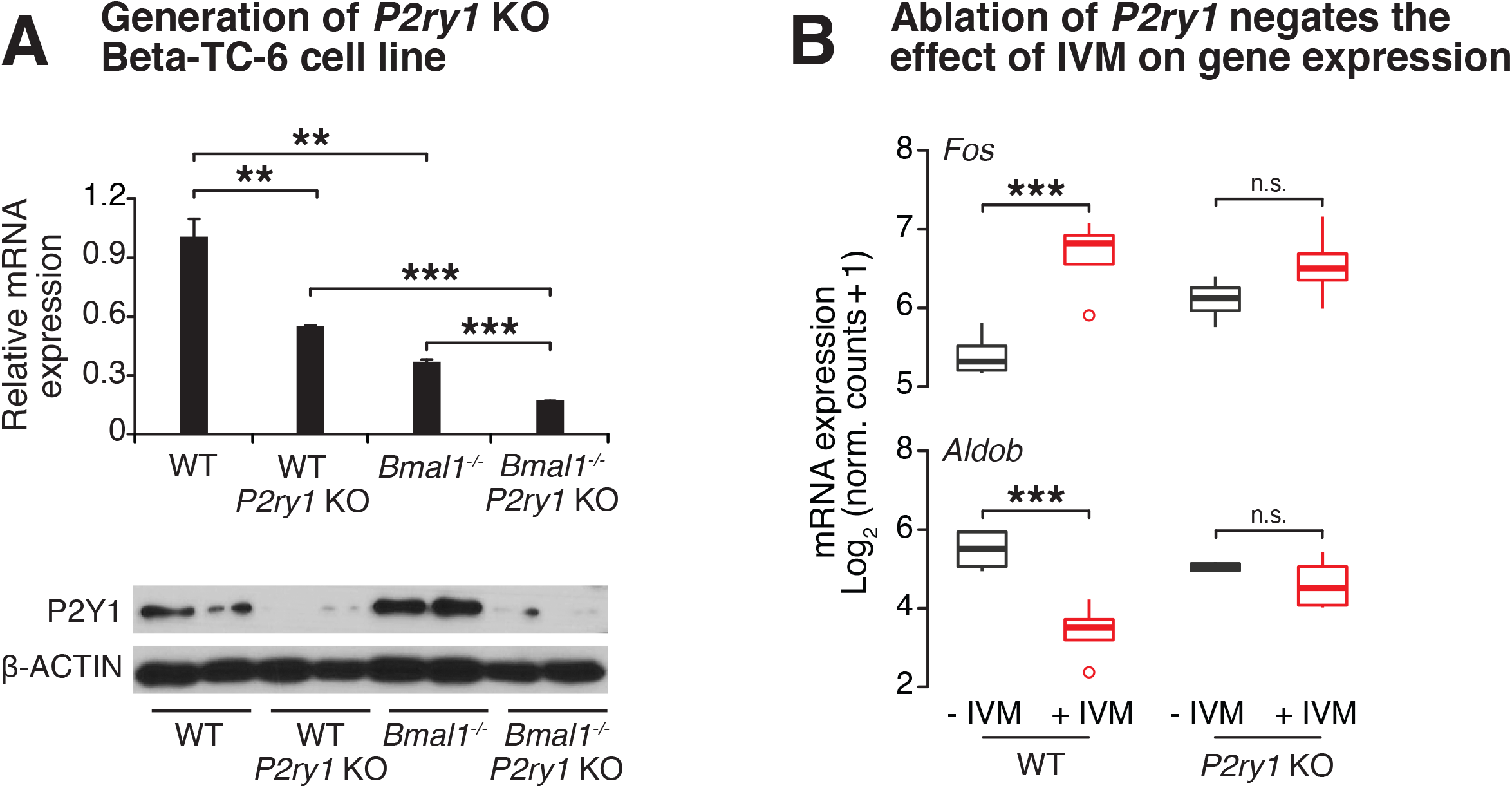
Genetic ablation of purinergic receptor P2Y1 in Beta-TC-6 cells blunts effect of IVM on gene expression. **(A)** Quantitative real-time PCR screening for disruption of *P2ry1* gene expression (n=3-4/genotype) (*top*). Decreased P2Y1 receptor protein expression by Western blot in WT and *Bmal1*^*-/-*^ Beta-TC-6 cells after genetic disruption (*bottom*). **(B)** Loss of effect of IVM on gene expression in *P2ry1* mutant β cells identified by RNA-sequencing (n=4/genotype/condition). Dots represent values that exceed 1.5-fold of the interquartile range. All values represent mean ± SEM. * p<0.05, ** p<0.01, *** p<0.001.

